# Circularization of Single-Stranded DNA Donor Template Unleashes the Power of Non-Viral Gene Delivery for Long-Term HSCs editing

**DOI:** 10.1101/2025.02.19.638978

**Authors:** Gil Letort, Aymeric Duclert, Diane Le Clerre, Isabelle Chion-Sotinel, Roger Salvatori, Emilie Dessez, Margaux Sevin, Marco Rotondi, Cosimo Ducani, Philippe Duchateau, Julien Valton

## Abstract

Over the past decade, non-viral DNA template delivery has been used with engineered nucleases to target single-stranded DNA sequences in hematopoietic stem and progenitor cells. While effective for gene therapy, this method has been limited to short DNA donor templates, restricting its applications to gene corrections. To expand its scope, we developed an editing process using kilobase-long circular single-stranded DNA donor templates and TALEN technology. Our results show that the CssDNA editing process achieves high gene insertion frequency in viable HSPCs. Compared to a conventional AAV-editing process, CssDNA-edited HSPCs show a higher propensity to engraft and maintain gene edits in a murine model. This positive outcome is partly due to higher levels of primitive edited HSPCs, a more quiescent metabolic state, and elevated expression of bone marrow niche adhesion markers. Our findings highlight the strong potential of CssDNA as a universal and efficient non-viral DNA template for gene therapy applications.

## Introduction

Gene therapy strategies utilizing hematopoietic stem and progenitor cells (HSPCs) hold the promise of delivering a lifelong supply of therapeutic agents to patients. Gene editing techniques expand these therapeutic prospects by employing engineered nucleases to introduce sequence-specific double-strand breaks (DSBs) and promote the targeted insertion of therapeutic genes vectorized as DNA donor templates. While viral vectors, particularly Adeno-associated viruses (AAVs), are the most prevalent and efficient carriers of DNA donor templates *in vitro*^1–6^, they have been recently associated with genotoxicity, impairment of HSPCs engraftment in murine model as well as loss of gene insertion events^7–12^. In this context, non-viral DNA templates recently emerged as promising alternatives to AAV6. Non-viral DNA templates come in different formats. The first format explored was linear double-stranded DNA (LdsDNA). When vectorized with engineered nuclease, LdsDNA enables gene correction and gene insertion in multiple cell types. But its recognition by intracellular DNA sensing pathways triggers cellular toxicity and impairs the overall efficiency of gene editing, especially in HSPCs^13,14^. To overcome this hurdle, linear single-stranded DNA (LssDNA) was considered. LssDNA, mostly used as short linear oligonucleotides (LssODNs, <300b), promotes efficient gene correction in Long-Term Hematopoietic Stem cells (LT-HSCs) without apparent cellular toxicity^8,9,11,15^. This correlated with an efficient engraftment of edited HSPCs in murine model and to the maintenance of their corrective events. Despite this, the limitation of short LssODNs to single point mutation corrections led the field to explore longer LssDNA templates (>1Kb) for full gene insertion in primary cells. Although highly efficient in T-cells, LssDNA showed a dose-dependent cellular toxicity and displayed low levels of gene insertion efficacy in HSPCs^16^, likely due to its degradation by exonucleases in the cytoplasm^17^.

Circular single-stranded DNA (CssDNA) offers a promising solution to the aforementioned challenges observed with different DNA donor template formats. Indeed, it may mitigate the activation of intracellular DNA sensing pathways that evolved to specifically recognize foreign dsDNA and is inherently resistant to exonuclease attacks. Recent proof-of-concept studies on CRISPR-CAS9/CssDNA-mediated gene insertion have demonstrated up to 25% gene insertion at multiple endogenous loci^18^. These promising results were very recently confirmed in HSPCs although the editing frequency and viability obtained seemed suboptimal (∼60 % cell viability and ∼20 % KI efficiency). Most importantly, the functionality of the resulting edited-HSPCs was not assessed *in vitro* and *in vivo*^19^. Furthermore, there has been no direct *in vitro* and *in vivo* comparison with AAV6, the standard DNA template vectorization method for targeted gene insertion. Therefore, a thorough investigation of CssDNA’s potential for gene insertion in HSPCs and comparison with other DNA vectorization methods, including linear single-stranded DNA (LssDNA) and AAV6, is essential for gauging its absolute and relative potential for the field of cell and gene therapy.

Here we developed a gene insertion process incorporating non-viral DNA donor templates (LssDNA or CssDNA) and TALEN technology for HSPCs editing. Our findings indicate that the CssDNA editing process yields a 3-to 5-fold higher gene Knock-in (KI) frequency than LssDNA process, with efficiencies surpassing 40%. Notably, this correlates with increased cell viability and a reduction in the frequency of gene knock-out (KO, via the promotion of insertions and deletions) in CssDNA-edited cells compared to those edited with LssDNA. These outcomes are consistent across various DNA donor template lengths (from 0.6 to 2.2 kb) inserted at multiple loci and are transportable to primary T-cells. Further comparative analysis of CssDNA and AAV gene insertion processes using a combination of Flow cytometry, CITE-Seq transcriptomics and long-term engraftment in NCG mice, suggest the superiority of CssDNA over AAV6 gene editing process. Our data also unveiled key transcriptomic and phenotypic distinctions associated to the efficacy of both gene insertion processes, including Long-Term HSC enriched subpopulation (LT-HSCe) number, editing frequency, and bioenergetic status. This work underscores the strong potential of combining CssDNA delivery with engineered nuclease for gene insertion in Long-Term repopulating HSCs and marks a crucial step towards advancing next-generation cell and gene therapies.

## Results

### Circular single-stranded DNA molecules can be used as DNA donor template for efficient TALEN-mediated gene insertion in HSPCs

To assess the potential of ssDNA molecule and TALEN editing to promote gene insertion in HSPCs, we chose the exon 1 of the *B2M* gene as endogenous genomic target. We used a *B2M* exon 1-specific TALEN (TALEN_B2M_) and designed promoterless DNA templates encoding cell surface reporters (CSR_1_ and CSR_2_) flanked by 300 bases homology arms specific for the *B2M* locus. These specific template designs allowed to seamlessly rewire the *B2M* gene in a disruptive or non-disruptive manner and to re-express the CSRs (Figure 1a and supplementary figure 1a). Because *B2M* is uniformly expressed in HSPCs and all their progenitors, this experimental system allows high-content, multiparametric single-cell level characterization of edited HSPCs and progenitor cells using either flow cytometry (FCM) and/or CITE-Seq analysis.

**Figure 1.**
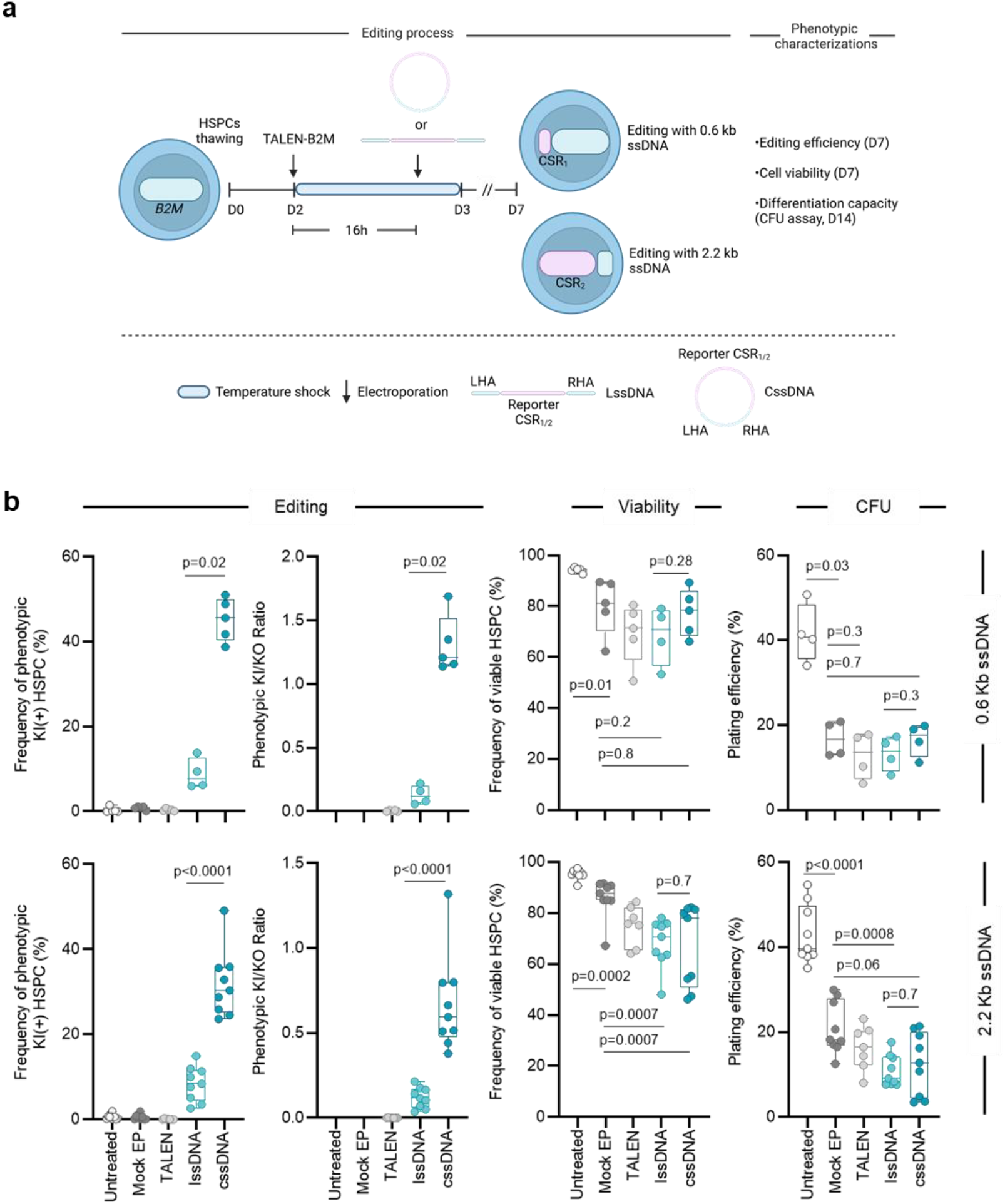
Circularization of ssDNA increases the overall efficiency of TALEN-mediated Knock-in in HSPCs. **a**. Representative schema of HSPCs editing protocol using an mRNA encoded TALEN targeting the *B2M* locus and LssDNA or CssDNA as DNA donor templates to insert a tag (0.6 kb) or a reported gene (2.2 kb) via non-disruptive and disruptive insertions, respectively. mRNAs encoding a viability enhancer and a HDR enhancer (Via-Enh01 and HDR-Enh01, respectively) were also incorporated in the first transfection. The timing is indicated in days (D0-D7). Edited HSPCs obtained 7 days post thawing were characterized by flow cytometry to assess the level of phenotypic knock-in (KI) of DNA donor templates and phenotypic knock-out (KO) of *B2M* as well as their viability. Their differentiation capacity into erythroid and myeloid progenitors was also assessed by colony forming unit (CFU) assay. **b**. Flow cytometry results illustrating the frequency of cells harboring KI events, the ratio KI/KO, the viability and plating efficiency of HSPCs either untreated, electroporated (Mock EP), edited with TALEN only (TALEN), or edited with TALEN and LssDNA or CssDNA donor templates (LssDNA or CssDNA, respectively). **Top and bottom panels** illustrate results obtained with 0.6 kb and 2.2 kb DNA donor templates, respectively. On each box plot, the central mark indicates the median, the bottom and top edges of the box indicate the interquartile range (IQR), and the whiskers represent the maximum and minimum data point. Each dot represents data obtained from one HSPCs donor.

Based on our former work^11^, we set up a 4 days-long gene insertion process consisting in HSPCs thawing (D0), followed by an expansion phase (D0-D2), a transfection of 3 mRNA molecules encoding the TALEN_B2M_, the HDR-Enh01 and the Via-Enh01^11^ (D2) followed by a second transfection (D3) of the ssDNA donor template. These steps were followed by one (D4) or multiple days of culture (D7 or D14) depending on the read out to be performed (figure 1a). Using this experimental setup, we first assessed the influence of the format of ssDNA donor template (Linear or Circular) on the frequency of gene editing (phenotypic gene knock-in, KI, and gene knock-out, KO) as well as on the viability and differentiation capacity of edited HSPCs (Figure 1 and supplementary figure 1). Three negative controls, consisting in untreated cells and electroporated cells with and without TALEN (“Untreated”, “TALEN” and “Mock EP” groups, respectively) were performed as references. Our results obtained with the 0.6kb-long ssDNA molecule encoding the CSR_1_ showed that both ssDNA formats elicited precise and non-disruptive KI at the *B2M* locus as demonstrated by the detection of a CSR_1_(+) HSPCs population by FCM (supplementary figure 1b and c). Interestingly, CssDNA promoted 5-fold higher KI than LssDNA, (Figure 1b top panel, 45.2% ± 5.0% and 8.8% ±3.6%, mean ± SD, respectively and supplementary figure 1c) with up to 51% of KI. This correlated with a 10-fold higher KI/KO ratio obtained with the CssDNA compared to the LssDNA (Figure 1b top panel, 1.30 ±0.22 and 0.13 ±0.07, mean ratio ± SD, respectively), indicating that CssDNA was more efficiently integrated at the *B2M* locus than the LssDNA. Although the electroporation process significantly impacted HSPCs viability with respect to untreated cells (Figure 1b top panel, median cell viability 81.3% and 94.5%, respectively, p-value=0.01), addition of TALEN_B2M_, LssDNA or CssDNA did not further impact it significantly (Figure 1b top panel, median cell viability 71%, 71% and 78%, respectively). This indicated that the electroporation process was mainly responsible for the slight loss of HSPC viability observed after gene editing. A similar pattern was observed by colony forming unit (CFU) assay. Indeed, HSPC electroporated alone or edited in the presence of TALEN_B2M_, LssDNA or CssDNA were able to differentiate in a similar fashion (Figure 1b top panel, median plating efficiency of 17%, 14%, 14% and 17%, respectively).

To assess if our editing process could enable longer gene insertion in HSPCs, we performed similar experiments with a 2.2 kb ssDNA template encoding CSR_2_, designed for disruptive insertion at the *B2M* locus. Our results showed that both ssDNA formats elicited precise and disruptive KI at the *B2M* locus as demonstrated by the detection of a CSR_2_(+) HSPCs population by FCM. CssDNA promoted higher KI and KI/KO ratio than LssDNA (figure 1b, bottom panel, 3.6-fold and 5.5-fold respectively, supplementary figure 1b and c) with up to 49% of KI. HSPC viability and differentiation capacity observed after 2.2kb ssDNA editing followed the same trend as the one observed with the 0.6kb ssDNA. Together these results indicate that CssDNA and TALEN_B2M_ editing process promoted significantly higher gene insertion than LssDNA in HSPCs without severely impacting HSPCs viability and differentiation capacity.

To gauge the transposability of CssDNA and TALEN editing for gene insertion at other loci in HSPCs, we designed 3 additional cssDNA donor templates specific for AAVS1, CD11B and S100A9 loci, demonstrated earlier to allow pan-lineage or myeloid-specific expression of therapeutic transgenes^20^ (supplementary figure 2). Using the aforementioned gene editing protocol, loci-specific TALEN and promoterless matrices encoding the CSR_3_, we showed that AAVS1, CD11B and S100A9 loci were efficiently modified, reaching similar gene insertion frequency and viability than the ones obtained with TALEN_B2M_ and CssDNA encoding CSR_2_ (supplementary figure 2b, 34% ± 9.6%, 26% ± 6.6% and 39% ± 10.5%, mean KI frequency ± SD, respectively). This additional dataset indicates that CssDNA could be used to promote efficient gene insertion at multiple loci in HSPCs.

To verify the suitability of CssDNA for HSPC-based ex-vivo gene therapy applications, we investigated whether the CssDNA editing process allowed gene insertion in Long-Term repopulating Hematopoietic Stem Cells (LT-HSC). For that purpose, we assessed the ability of HSPCs edited by the TALEN_B2M_ locus and the CssDNA encoding CSR_2_, to engraft in an NCG murine model, to differentiate and maintain their gene insertion events, 16 weeks post injection onset (figure 2a). A control experiment was performed with HSPCs edited with an adeno-associated virus editing process (AAV process, TALEN_B2M_ and AAV6 encoding the CSR_2_), since AAV is considered as the conventional and therapeutically relevant DNA donor template vectorization method for targeted gene insertion in human HSPCs^1–6^. To allow for rigorous side-by-side comparisons of both processes and mitigate the occurrence of the confounding side effects usually associated to AAV6^12,21^, we defined the minimal MOI of AAV6 (350 vg/cells) to reach similar gene insertion efficiency than the one obtained with the CssDNA editing process. By design, both CssDNA and AAV editing processes elicited similar KI frequency (supplementary figure 3a left panel, 30.7% ± 2.1% and 26.1% ± 5.4%, mean ± SD, respectively), KI/KO ratio (supplementary figure 3a left panel, 0.61 ± 0.08 and 0.53 ± 0.14, mean ± SD, respectively) and cell viability (supplementary figure 3a middle panel, 81.0% ± 1.0% and 82.5% ± 4.3%, mean ± SD, respectively) using 3 HSPC donors. As demonstrated earlier, edited HSPCs were able to differentiate *in vitro* as observed by CFU assay (supplementary figure 3a, right panel). Interestingly, although not significant, the AAV editing process led to a 1.7-Fold increase of plating efficiency compared to its CssDNA counterparts (supplementary figure 3a right panel, 31.3% ± 5.2% and 19.2% ± 4.5%, mean ± SD, respectively).

**Figure 2.**
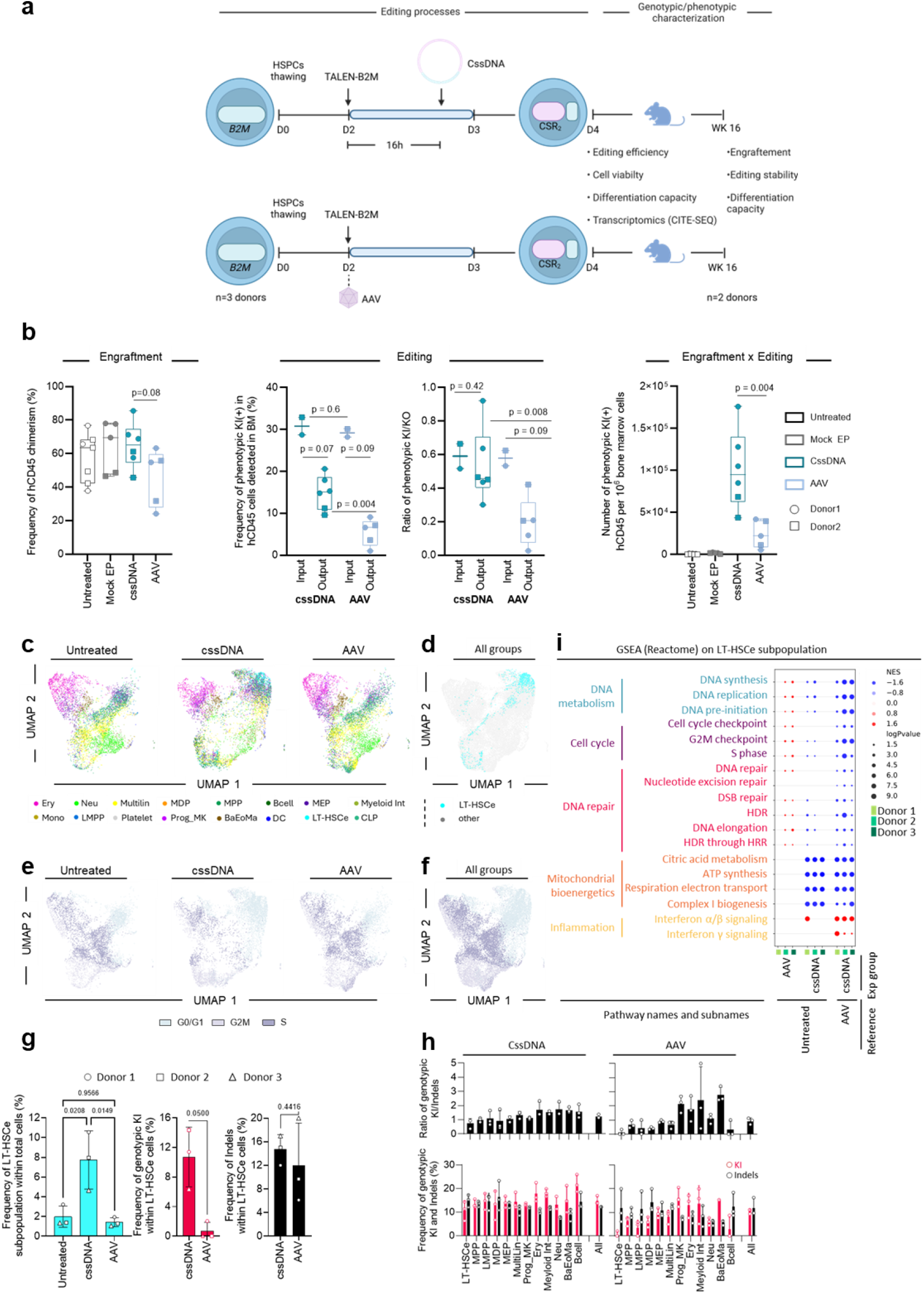
CssDNA/TALEN-mediated knock-in process leads to a higher engraftment of KI(+) HSPCs compared to the AAV/TALEN reference process. **a**. Representative schema of HSPCs editing protocol using an mRNA encoded TALEN targeting the *B2M* locus and cssDNA or AAV (MOI=350 vg/cell) as DNA donor templates to insert a reported gene (CSR_2_, 2.2 kb) via disruptive insertion. mRNAs encoding a viability enhancer and a HDR enhancer (Via-Enh01 and HDR-Enh01, respectively) were also incorporated in the first transfection. The timing is indicated in days (D0-D7). Edited HSPCs retrieved 7 days post thawing (D7) were characterized by flow cytometry to assess the level of knock-in (KI) of DNA donor templates and knock-out (KO) of *B2M* as well as their viability. Their differentiation capacity into erythroid and myeloid progenitors as well as their transcriptomics profile were also assessed by colony forming unit (CFU) assay (supplementary figure 4b) and CITE-seq, respectively. Edited HSPCs retrieved 4 days post thawing (D4), were also injected in NCG mice to assess their ability to engraft, differentiate and keep their editing events, 16 weeks after injection onset. **b**. In vivo experimental results illustrating the level of human CD45+ cells (hCD45) engraftment and of KI frequencies and KI/KO ratio determined either before mice injection (input), or in hCD45+ cells engrafted in the bone marrow (BM) of NCG mice, 16 weeks after cells injection onset (output). Two-way ANOVA followed by Bonferroni multi-comparison test. P-values are indicated. The product of the frequency of hCD45+ cells engraftment and frequency of KI is also shown to illustrate the overall efficiency of each HSPC editing process. Mann–Whitney two-tailed non-parametric unpaired test with a confidence interval of 95%. P-value is indicated. On each box plot, the central mark indicates the median, the bottom and top edges of the box indicate the interquartile range (IQR), and the whiskers represent the maximum and minimum data point. Each dot represents data obtained from one HSPCs donor. **c**. UMAP plots showing aggregated 5’scRNA CITE-Seq data obtained from HSPCs either untreated, edited with TALEN and AAV or cssDNA donor templates (AAV or cssDNA respectively), 4 days post thawing (D4) and at the time of NCG mice injection onset (n = 3 independent biological donors). The different cell subpopulations identified are illustrated by a color code indicated at the bottom of the graph. The Long-term HSC-enriched cell subpopulation is indicated as LT-HSCe and definition of each subpopulation is documented in the methods section **d**. UMAP plots aggregated from 5’scRNA CITE-Seq data obtained from all experimental groups showing the position of LTHSCe subpopulation. **e and f**. UMAP plots showing the cell cycle phases (G0/G1, G2M and S) of each cell identified in each experimental groups and in all groups, respectively. **g. left panel**, illustrate the frequency of LT-HSCe within all subpopulations. Two-way ANOVA with Bonferroni post-tests. P-values are indicated. **g middle and right panels**, illustrate the frequency of KI and KO within the LT-HSCe subpopulation, respectively. Paired t-test. P-values are indicated. Each dot represents data obtained from one HSPCs donor. **h**. plot showing the frequency of KI(+) and KO(+) in each subpopulation found in the CssDNA and AAV-edited HSPCs (n=3 donors aggregated, subpopulations identified with fewer than 100 cells are not displayed). **i**. Gene Set Enrichment Analysis (GSEA) obtained in LT-HSCe to compare the AAV and CssDNA experimental groups to the untreated reference group and to directly compare the cssDNA group to the AAV reference group. Normalized Enrichment Score (NES) as well as Log P value obtained for each donor (n=3 independent HSPC donors) are illustrated by a red/white/blue color code and size of the dots, respectively. The different pathways found to be significantly up (red) or down (blue) regulated in the different experimental group comparisons are identified as individual pathways and aggregated in subgroups for the sake of clarity.

HSPCs donors 1 and 2, showing the most similar editing outcomes *in vitro* between CssDNA and AAV6 processes, were used for further side-by-side comparison *in vivo*. Our results obtained 16 weeks post injection onset, showed that both processes led to similar HSPC engraftment frequencies, although HSPCs edited by the AAV process showed the lowest one (figure 2b left panel, 65.0% ± 13.6 and 46.0% ± 17.0%, mean ± SD, for CssDNA and AAV6 processes, respectively). They also elicited similar levels of HSPC differentiation into Lymphocyte T and B, myeloid cells, monocyte, macrophages and neutrophiles with respect to untreated cells (supplementary figure 3b). Notably, the CssDNA editing process allowed for preservation of significantly higher number of KI events than the AAV editing process. Indeed, we found a 3-fold higher KI frequency in hCD45 cells engrafted in animal injected with CssDNA-edited HSPCs compared to those injected with AAV-edited HSPCs (figure 2b middle panel, 14.9% ± 4.1% and 5.5% ± 3.2% mean ± SD, respectively). This trend was consistent among the different cell subpopulations studied (supplementary figure 3c). In addition, the CssDNA editing process allowed to keep a KI/KO ratio constant between the input and output, in contrast with the drop observed with the AAV6 editing process (Figure 2b middle panel, 0.59 versus 0.53, and 0.58 versus 0.18, mean KI/KO ratio obtained at input and output for CssDNA and AAV processes, respectively). Ultimately, the overall engraftment efficiency of edited HSPCs, calculated by multiplying the bone marrow engraftment and output KI frequencies, was 5-fold higher for HSPCs edited using the CssDNA process compared to those edited using the AAV process (figure 2b right panel, 9.20E4 versus 1.85E4, mean number of KI(+) HSPCs per 1E6 bone marrow cells). Collectively, our findings indicate that the CssDNA editing process enables efficient targeted gene insertion in HSPCs, producing cells capable of engrafting in the bone marrow of the NCG murine model at significantly higher levels than those edited with the AAV6 process.

To understand the significant differences observed between CssDNA- and AAV6-edited HSPCs engraftment in murine model, an in-depth characterization of cells gathered right before their injection in NCG mice (D4), was performed using single cell CITE-Seq analysis. A total of 22359 encapsulated cells coming from 3 different donors (including those used for *in vivo* engraftment analysis), enabled us to analyze a median of 2414 cells per experimental group and per donor, with a median of 6318 genes analyzed per cells (table 1). This comprehensive dataset enabled us to first decipher the different cell populations present in the cellular mixture using as reference, the comprehensive dataset documented in^22^ (Figure 2c and supplementary figure 4). Among them, we identified the most primitive subpopulation harboring, among others, the markers commonly expressed by Long-Term HSCs (*AVP, HLF, ITGA3, CRHBP, THY1* and *PROM1*, Figure 2d and supplementary figure 4)^11,22–24^. This subpopulation, displaying a majority of cells in G0/G1 (figure 2e,f and supplementary figure 4), will be called Long-Term HSC enriched subpopulation (LT-HSCe) in the following, for the sake of simplicity.

Interestingly, the frequency of LT-HSCe was found to be 5-fold higher in the CssDNA edited HSPCs compared to the AAV-edited or untreated HSPC groups (Figure 2g left panel, 7.7% ± 3.0%, 1.4% ± 0,5%, and 2.0% ± 1%, median ± SD, respectively). Further analysis of the sequence of B2M transcripts as well as surface expression of the CSR_2_, enabled us to confidently identify cells displaying phenotypic and genotypic KI as well as genotypic KO, referred in the following as insertion and deletion (indels) events, within each subpopulation of edited HSPCs (supplementary figure 5a). Using this additional layer of information, we determined that the frequency of LT-HSCe harboring KI events (KI(+) LT-HSCe) obtained using the CssDNA process was 10-fold higher than that obtained using the AAV process (figure 2g middle panel, 10.7% ± 4.0% and 0.7% ± 1.2%, mean ± SD, respectively). In stark contrast, the frequency of LT-HSCe harboring indels events (Indels(+) LT-HSCe) were surprisingly similar from one process to another (figure 2g right panel, 14.7% ± 2.5% and 12.0% ± 7.2%, mean ± SD, respectively). This phenomenon was also observed in other most primitive subpopulations including Late Multipotent Progenitor and Macrophage Dendritic Progenitor (LMPP and MDP, respectively) but not in differentiated subpopulations including Progenitor Megakaryocytes, Erythroid and Myeloid intermediates (Prog_MK, Ery, Myeloid Int, respectively) as non-limiting examples, figure 2h). Deeper investigation of the KI events within each cell cycle phase of LT-HSCe subpopulation, showed that CssDNA-edited KI(+) LT-HSCe were mainly set in G0/G1 phase (80%, Supplementary Figure 5b, left panel). Unfortunately, such analysis could not be rigorously performed on AAV-edited LT-HSCe due to the low number of KI(+) LT-HSCe cells identified in this group (1 LT-HSCe KI(+) cell, Supplementary Figure 5b, right panel).

Nevertheless, we could compare the overall transcriptomics status of LT-HSCe detected in each experimental groups using Gene Set Enrichment Analysis (GSEA, Reactome, Figure 2i). Our results showed that pathways associated to DNA metabolism, cell cycle, DNA repair and mitochondrial bioenergetics were significantly downregulated in CssDNA-edited LT-HSCe compared to its AAV-edited and untreated counterparts. This indicated a lower overall proliferative status of the former group, in agreement with the conclusions made earlier from the cell cycle analysis and cellular fold expansion (Supplementary figure 3). Interestingly, the opposite phenomenon was observed with respect to inflammation pathways (IFNα/β and γ) indicating that CssDNA was more likely to be recognized by DNA sensing pathways than the AAV particles in the cytoplasm, in agreement a with former study^11^. Finally, we found that the CXCR4, CD44 and F11R, three surface markers involved in cellular adhesion to the bone marrow niche, were significantly upregulated in the LT-HSCe subpopulation of CssDNA-edited cells compared to their AAV-edited counterparts or untreated cells (Supplementary figure 5c, Log fold change of expression ≥1).

While CssDNA bears a strong potential for *ex vivo* HSPC-based gene therapy, it may also find applicability in other fields including CAR T-cell therapy and immuno-oncology. We thus also assessed the transposability of CssDNA and TALEN editing to T-cell engineering. To investigate that aspect, we established a T-cell-centric TALEN engineering protocol in the absence of Via-Enh01 and HDR-Enh01 encoding mRNA and used it to assess the efficiency of gene insertion mediated by 2.2 kb-long CssDNA and LssDNA templates encoding the CSR_2_ specific for the *B2M* locus (supplementary figure 6a). Our results show that CssDNA promoted higher frequency of gene insertion than the LssDNA, whatever the dose being used, with up to about 40% of KI (supplementary figure 6b and c). This process did not significantly impact the viability of edited T-cells at D7 (supplementary figure 6d, mean viable cell frequency >90% for both CssDNA-, LssDNA- and TALEN-edited T-cells) and allow them to proliferate, although to a lower extent than T-cells edited by TALEN_B2M_ alone (∼3.7-fold lower proliferation rate of CssDNA-treated T-cells compared to TALEN_B2M_ alone-treated T-cells, supplementary figure 6e).

## Discussion

The goal of this study was to evaluate the potential of linear and circular ssDNA donor templates (LssDNA and CssDNA) to enable TALEN-mediated gene insertion in HSPCs. Our findings demonstrate that both DNA donor templates enable precise gene insertion at endogenous loci. CssDNA significantly and markedly increases the efficiency of gene insertion frequency by a up to 5-fold compared to LssDNA, with absolute values surpassing 40% using a 2.2kb-long ssDNA template. This unprecedent efficiency of CssDNA insertion correlates with a significant decrease of Indels events, considered as by-products of the gene insertion process. Our CssDNA-editing process can be used to insert kilobase-long genes at multiple independent loci and is transportable to other cell types, including primary T-cells used for therapeutic applications. Comparative analysis with a conventional AAV-editing process reveals that HSPCs edited by CssDNA display a higher propensity to engraft and maintain their gene insertion edits in a murine model compared to HSPCs edited by AAV. This difference could be explained in part, by a significantly higher levels of KI(+) LT-HSCe subpopulation, a more quiescent metabolic state and an elevated expression of bone marrow niche adhesion markers, in CssDNA-compared to AAV6-edited HSPCs. Our findings underscore the strong potential of CssDNA as a universal and efficient non-viral DNA template for cell and gene therapy applications.

We found that the efficiency of TALEN-mediated gene insertion was up to 5-fold higher with CssDNA than LssDNA donor template. This significant difference, observed in HSPCs and primary T-cells, using two lengths of ssDNA templates and across several donors, is consistent with results obtained in K562 and HEK293T cell lines edited with CRISPR-Cas9 nuclease^18^. It also aligned with a very recent study published just before the submission of this work, showing that circularization of ssDNA increases gene insertion efficiency by 5-fold in CRISPR-CAS9 edited K562 cells^19^. In this study, CssDNA was also used as DNA donor template in combination with CRISPR-CAS9 nuclease or nickase (vectorized as RNP or mRNA) to insert, in a targeted fashion, kilobase-long DNA in primary T-cells with KI efficiency ranging from 15% to 40% and a cell viability around 80% (depending on the locus edited and T-cell donor). Targeted insertion of CssDNA was also demonstrated in other cell types with variable cell viability and KI efficiency outcomes including in iPSCs (∼30% cell viability and 40-60% KI efficiency), NK cells (∼80% cell viability and 20-40% KI efficiency), B-cells (∼80% cell viability and 25% KI efficiency), and HSPCs (∼60% cell viability and ∼20% KI efficiency). Our work thus corroborates and extends these former investigations by demonstrating that circularization of ssDNA improves KI efficiency with up to 49% of multi kilobase-long gene insertion efficiency and a median viability of about 80% in HSPCs (figure 1). It also showed that multiple engineered nuclease platforms, including CRISPR-CAS9 and TALEN, could be used alongside CssDNA to promote efficient, non-viral gene insertion in different primary cells.

The difference in gene insertion efficiency observed in the presence of LssDNA and CssDNA could be due to multiple factors. One factor could be the hydrodynamic volume and shape of ssDNA, which may affect its penetration of the nuclear membrane during electroporation. Another more likely factor, could be their stability and susceptibility to Trex1 exonuclease-dependent hydrolysis, as recently demonstrated in the context of CRISPR-CAS9-mediated insertion of short LssODN^17^. Adding phosphorothioate bonds at the 3’ end of LssDNA could be beneficial by preventing its hydrolysis, increasing its stability and by extension, its insertion efficiency^17,25^. Although the incorporation of such modifications in long LssDNA molecules is not streamlined yet, further research should be conducted to investigate this avenue.

Nevertheless, we show that adapting the format of ssDNA donor template through simple circularization is sufficient to significantly improve the efficacy of gene insertion in primary cells. This improvement was demonstrated by rigorously comparing LssDNA- and CssDNA-mediated gene insertion in the presence of an mRNA encoding HDR-Enh01, a genetically encoded inhibitor of 53BP1, known to increase the efficiency of gene correction in HSPCs without generating additional and detectable adverse events^11^. The systematic incorporation of HDR-Enh01 in our gene editing process thus participates to the high levels of gene insertion obtained in this study. Nonetheless, achieving these high levels of gene insertion did not require Cas9 targeting sequences (CTS)^26^ or HDR-boosting modular sequences^27^ to reach high levels of gene insertion, although their incorporation in CssDNA could be beneficial. In addition, it did not need small molecule-based inhibition of DNA-PK and Polθ, an elegant approach known to dramatically increase the efficiency of gene insertion/correction in multiple cell types^28,29^ but is likely to promote large scale genomic alterations in HSPCs when used in the presence of short ssDNA donor templates^30^.

Our CssDNA edited process enables targeted insertion of multi kilobase-long gene in HSPCs, achieving unprecedented efficiency with respect to recent studies^16,31^. However, beyond editing efficiency, the functionality of edited cells is of paramount importance for their effective translation into HSPC-based gene therapy products. In this regard, the choice of the DNA donor template format as well as the editing process appears to be critical^8,9,11^. With these considerations in mind, we assessed the functionality of CssDNA-edited HSPCs *in vivo* using an NCG murine model and analyzed their transcriptomic status at a single cell level. This in-depth characterization was performed alongside AAV-edited HSPCs, since AAV is regarded as a reference of DNA donor template format for ex vivo HSPCs-based gene therapy applications^1–6^ and was the first to be used in the clinic in the context of sickle cell disease gene correction therapy (CEDAR trial, NCT04819841). Our results demonstrate that both groups of edited HSPCs were able to engraft and differentiate in NCG mice, indicating they contain a fraction of Long-Term repopulating HSCs. However, the level of engrafted HSPCs harboring gene insertion events (KI(+) HSPCs) was 5-fold higher in animals injected with CssDNA-edited HSPCs compared to those injected with AAV-edited HSPCs (figure 2b right panel). This differential outcome was obtained despite similar levels of KI observed at input for both groups and the extremely low AAV6 MOI used to mitigate the toxicity of this DNA vectorization approach in HSPCs (supplementary figure 3a left panel)^12,21^. These results are reminiscent of former works^8,9,11^ and suggest that the CssDNA editing process enables the generation of higher levels of Long-Term repopulating HSCs harboring KI events compared to its AAV counterpart. In agreement with this hypothesis, transcriptomic analysis performed on HSPCs just before their injection in mice showed significantly higher numbers of LT-HSCe in CssDNA-edited HSPCs than in AAV-edited HSPCs (figure 2g, left panel, 5-fold). In addition, CssDNA-edited LT-HSCe showed a significantly higher frequency of KI and a higher expression of bone marrow niche adhesion markers than their AAV counterparts (figure 2g middle panel, 10-fold and supplementary figure 5c, > 1 log fold change of CXCR4, CD47 and F11R expression, respectively). We hypothesize that the combination of these different features participates to an efficient engraftment of CssDNA-edited LT-HSCe and to the maintenance of their KI event in our murine model.

Interestingly, while the phenotypic KI/KO ratio remained stable after long term engraftment of CssDNA-edited HSPCs (figure 2b, middle panel), it significantly decreased in their AAV-edited counterparts. This suggests that the CssDNA editing process generates similar levels of phenotypic KI and KO editing events within Long-Term repopulating HSCs, whereas its AAV counterpart may favors phenotypic KO over KI events in Long-Term repopulating HSCs. In accordance with this finding, our transcriptomic analysis showed that while the CssDNA-editing process elicited similar level of genotypic KI and indels events within LT-HSCe, the AAV-editing process rather favored genotypic indels over KI events (Figure 2g and h). This imbalance of KI over indels events was expected in AAV-edited LT-HSCe^7,21,32,33^. Indeed, genotypic KI events mainly occur in the S/G2M phases through the homology-directed repair (HDR) pathway. These events are thus less favorable in quiescent and primitive cells compared to indels events which can happen in any cell cycle phases through the non-homologous end joining (NHEJ) pathway^7,21,32,33^. In agreement with this, such imbalance was also observed in other primitive subpopulations but was found normalized (KI/indels ratio ∼1), if not inverted (KI/indels ratio >1), in most of the differentiated subpopulations (figure 2g and h). Of note, this imbalance can also be due to the higher toxicity of AAV particles on primitive subpopulations compared to differentiated subpopulations and further work is now needed to fully address this point.

Surprisingly, in contrast to the observation described above for the AAV-editing process, the CssDNA-editing process elicits similar levels of KI and indels events in the most primitive subpopulations including MPP, LMPP, MDP and LT-HSCe (figure 2g and h). Moreover, it was also found to skew the KI events toward the LT-HSCe set in G0/G1 phases over those set in G2M or S phases. This pattern could be due to multiple parameters including the overall engineering process and/or to an alternative mechanism of DNA template insertion as described earlier for LssODN-mediated gene correction^34^. Further work is now needed to fully address this point, which lies outside the scope of this study.

The success of HSPC-based ex vivo gene therapy relies on multiple factors ranging from the efficiency of gene modification/insertions, the stemness of edited HSPC subpopulation, the viability, differentiation capacity and long-term engraftment potential of the resulting edited HSPCs. Multiple avenues have been explored in the past decade to improve the efficiency of targeted gene insertion in mobilized-HSPCs, a cell population relevant for ex vivo gene therapy. Utilization of engineered nuclease and AAV6 as DNA donor template showed highly efficient gene targeting outcomes without affecting the overall HSPCs viability and differentiation capacity *in vitro*. These outcomes were obtained with different engineered nuclease platforms, were reproducible across multiple loci and obtained by several teams across the world^1–6^. The vivid enthusiasm for this approach was however dampened by the suboptimal functionality of edited HSPCs including their engraftment capacity in murine model as well as their poor ability to maintain their gene insertion edits after long-term engraftment *in vivo*^7–9^. Further in-depth investigations shed light onto these phenomenon by demonstrating the propensity of AAV6 to activate P53 pathways in a dose dependent manner, and reduce the clonal repertoire of engrafted HSPCs^10–12^.

These converging evidences led the field to explore other strategies, including those aimed at reducing the confounding toxicity of AAV6^21^ and those using alternative DNA donor template vectorization^8–11,15,35^. Our work further explores and expands the latter avenue by showing that CssDNA can be used as DNA donor template along with TALEN gene editing to efficiently promote non-viral gene insertion in LT-HSCs. We demonstrate that CssDNA-edited HSPCs are viable, functional and can engraft and differentiate in vivo in a xenograft murine model. We also demonstrate that it can be used to insert kilobase-long genes at multiple loci and in different cell types, including primary T-cells used for therapeutic applications. Finally, the CssDNA templates were synthesized using in vitro enzymatic DNA synthesis based on Rolling Circle Amplification^36^. This production process is highly scalable due to its reliance on isothermal enzymatic reactions. It is also potentially safer than fermentation-based DNA production methods^31,37^, which are often susceptible to endotoxin contamination. As a result, CssDNA production can be seamlessly implemented as a GMP-compliant process. We thus believe that CssDNA bears strong potential and will be crucial in guiding the next generation of HSPCs-based gene therapies for the benefit of patients.

## IV-Methods

### CD34+ HSPCs sourcing and culture

Frozen CD34+ HSPCs purified from healthy donor G-CSF-mobilized and Plerixafor-mobilized peripheral blood were purchased from AllCells (Almeda). After thawing, CD34+ HSPCs were cultured at a concentration of 0.4×10^6^ cells/mL in complete medium: StemSpan II (Stemcell, #09655), 1X CD34 expansion supplement (Stemcell, #02691) and 1X penicillin-streptomycin (Gibco, #15140-122) at 37°C, 5% CO_2_. HSPCs were assessed for viability by Nucleocounter or by the expression of CD34 and viability marker by flow cytometry 2 or 5 days after gene editing. Flow cytometry staining was performed in Annexin V Binbing Buffer 1X (BD Pharmingen, #556454) with antibody panel documented in Supplementary Table 2. The gating strategy used to analyze cellular suspension is documented in supplementary supplementary figure 1.

### TALENs and DNA donor templates

Plasmids of the 4 TALEN arms, containing a T7 promoter and a polyA sequence (120 residues), were produced after assembly and linearized for mRNA *in vitro* transcription. TALEN mRNAs were produced using an in-house In vitro transcription (IVT) process.

For non-viral mediated gene insertion, a DNA donor template sequence containing the coding sequences of CSR1 (HA-Tag), CSR2 (HLAE trimeric construct^38^) or CSR3 (ΔLNGFR) flanked by 300-nt left and right homology arm sequences specific for the locus to modified (B2M, AAVS1, CD11b or S100A9) were designed. The corresponding LssDNA and CssDNA were produced by Moligo Technologies, using an enzymatic in vitro technology, using a proprietary protocol, partially based on the Rolling Circle Amplification approach documented in^36^. For viral-mediated gene insertion, the DNA donor template containing CSR_2_ and documented in^38^ was used to produce the corresponding AAV6 particles.

### HSPCs engineering

The procedures used to transfect and/or transduce HSPCs were adapted from our former work documented in ^11,39^. Briefly, two days after thawing, the cells were washed twice in BTXpress buffer and resuspended at a final concentration of 40×10^6^ cells/mL in the same solution. The cellular suspension (4×10^6^ cells) was mixed with 5 µg mRNA encoding each TALEN arm in the presence of 4 µg mRNA and 1µg mRNA encoding for HDR-Enh01 and Via-Enh01, respectively, in a final volume of 100 µl. The cellular suspension was transfected in 4-mm cuvette gap size using PulseAgile technology. The electroporation program consisted of two 0.1 ms pulses at 1000 V/cm followed by four 0.2 ms pulses at 130 V/cm. Immediately after electroporation, the HSPCs were transferred to a new plate containing prewarmed medium at a concentration of 2×10^6^ cells/mL and incubated for 15 minutes at 37°C.

For AAV-mediated transduction experiments, 15 minutes after TALEN electroporation, HSPCs were cultured at a concentration of 2×10^6^ cells/mL in the presence or absence of AAV6 particles (MOI = 350 viral genome/cell) and incubated for 15 minutes at 37°C. HSPCs were then incubated at 30°C overnight. For ssDNA transfection experiments, HSPCs were kept in culture at 30°C for 16 hours after TALEN electroporation, recovered and washed 2 times in PBS (Gibco, 10010056). A second electroporation was performed, using 1×10^6^ cells in presence or absence of 0.02 nmol of ssDNA in a final volume of 100 µl using Nucleofector 2b apparatus (Lonza, AAB-1001) and Human CD34+ Cell Nucleofector™ Kit (lonza, #VPA-10003) using U-08 program. After ssDNA transfection, HSPCs were seeded at a concentration of 2×10^6^ cells/mL and incubated at 30°C overnight. The following day, cells were seeded at a density of 0.3×10^6^ cells/mL in complete medium and cultured at 37 °C in the presence of 5% CO2.

### T-cells engineering

The procedure used to transfect T-cells were adapted from our former work documented in ^11,39^. Briefly, one day after thawing, T-cells were activated using Transact (Miltenyi#200-076-204, 60µL/ 10^6^ CD3(+) cells) and allowed to grow for 3 days. Activated T-cells were then recovered, washed twice in Cytoporation T buffer and resuspended at a final concentration of 50×10^6^ cells/mL in the same solution. The cellular suspension (5×10^6^ cells) was mixed with 5 µg mRNA encoding each TALEN_B2M_ arm in a final volume of 100 µl. The cellular suspension was transfected in 2-mm cuvette gap size using PulseAgile technology. The electroporation program consisted of two 0.1 ms pulses at 400 V/cm followed by four 0.2 ms pulses at 100 V/cm. Immediately after electroporation, the transfered were transferred to a new plate containing prewarmed medium at a concentration of 2×10^6^ cells/mL and incubated at 30°C. T-cells were kept in culture at 30°C for 6 hours after TALEN electroporation, recovered and washed 2 times in PBS (Gibco, 10010056). A second electroporation was performed using 2.5×10^6^ cells in absence or presence different quantities of ssDNA in a final volume of 100 µl using Nucleofector 2b apparatus (Lonza, AAB-1001) and Human T Cell Nucleofector™ Kit (lonza, #VPA-10002) using T-023 program. After ssDNA transfection, T-cells were seeded at a concentration of 2.5×10^6^ cells/mL and incubated at 30°C overnight. The following day, cells were transferred to 37 °C and cultivated in the presence of 5% CO2 for further assessment of gene editing efficiency and rate of proliferation.

### Colony forming unit (CFU) assays

The procedure used to perform CFU assay were adapted from our former work documented in^11^. Briefly, CD34+ HSPCs recovered two days after electroporation, were plated in methylcellulose (Stemcell, #04435) for Colony Forming Unit (CFU) assays. A total of 200-500 cells (CD34+) were resuspended in 100 µL of Stemspan II and transferred to an aliquot of 1 mL of methylcellulose, mixed, and plated in a Smartdish well (Stemcell, #27371). Cells were cultured for 12-14 days in methylcellulose according to the manufacturer’s instructions. At the end of the culture, colonies were automatically counted using a Stemvision (StemCell) automated colony counter to assess plating efficiency (number of colonies counted at day 14 / number of cells plated at day 0).

### Single-cell CITE-Seq procedure and bioinformatic analysis of sequencing results

Single-cell mRNA barcoding and library generation were performed following the 10X Genomics protocol from the Chromium Next GEM Single Cell 5’ Kit v2 (#1000263) as described in ^11^. Samples from different donors were multiplexed following a previously described cell-hashing protocol. Briefly, 200,000 edited or non-edited HSPCs (D4) were thawed and incubated with Human TruStain FcX™ Fc Blocking reagent (Biolegend, #422301) for 10 minutes at 4°C. Cells were stained with TotalSeq-C Hashtag antibodies diluted at 1/50 in Cell Staining buffer (Biolegend, #420201) for 30 minutes at 4°C. After washing, different conditions bearing different hashtags were pooled. 500,000 cells were stained for 30 min at 4°C with TotalSeq-C Universal Cocktail V1.0 (Biolegend, #399905) supplemented with 18 additionals TotalSeq-C antibodies at 0.25ug per 500,000 cells. The list of antibody-derived tag (ADT) used for CITE-Seq procedure is documented in supplementary Table 2. After washing cells were loaded into Chromium Single-Cell Chip (10X Genomics) at a target capture rate of ∼10,000 individual cells per sample. Gene Expression and Cell Surface libraries were checked using a Bioanalyzer High Sensitivity DNA kit (Agilent, #5067-4626) according to the manufacturer’s recommendations. Libraries generated were sequenced by Institut du Cerveau (ICM) IGenSeq plateformusing a NovaSeq X Plus sequencer from Illumina following 10X recommendations.

Raw demultiplexed sequences were processed by Cell Ranger multi v8 using human reference genome GRCh38 to generate the count matrices. Cells with less than 1000 genes or more than 9000 genes were filtered out, as well as cells with more than 10% mitochondrial genes. Data were processed using Seurat v5. Briefly, data were normalized, the 2000 most variable mRNA features were retained, before scaling and doing a Principal Component Analysis (PCA). Protein and mRNA features were integrated using Seurat WNN function FindMultiModalNeighbors. Cell cycle was predicted using CellCycleScoring function from Seurat. A dataset of CD34(+) high cells from 8 donors (about 190,000 cells) obtained from ^11^ was used as a reference to map our cells using Symphony^40^. Cell type acronyms were used in Figure 2 and supplementary figure 4 to name the different HSPCs subpopulations identified. We used LT-HSCe for Long-Term HSC enriched, the most primitive Hematopoietic Stem Cells identified in this dataset; MPP for Multipotent Progenitors; LMPP for Lymphoid Primed Multipotent Progenitors; CLP for Common Lymphoid Progenitors; MEP for Megakaryocytic Erythroid Progenitors; MDP for Monocyte-macrophage and Dendritic cells Progenitors; Prog MK for Progenitor Megakaryocytes; Ery for Erythroid Progenitors; Myeloid_Int for Myeloid Intermediate Progenitors ; Neu for Neutrophil Progenitors; Mono for Monocyte Progenitors; BaEoMa for Basophil Eosinophil Mast Progenitors; Prog DC for Progenitor Dendritic Cells and Bcells for Progenitors and Precursor B cells. KI resulting from AAV or CssDNA produced a large increase of the expression of HLA-E at the surface which could be detected by one of the antibodies present in the CITE-seq. Taking as a reference the untreated samples, a threshold on HLA-E ADT signal was determined above which a cell could safely (false positive rate estimated at 1%) be evaluated as having KI from its phenotype (called phenotypic KI). Differential gene expression was assessed using Seurat FindMarkers with default parameters but setting no minimum for the log fold change and min.pct. Gene Set Enrichment Analysis (GSEA) was carried out using using Reactome pathways retrieved from MSigDB with the R package msigdbr. Sankey diagrams were generated using Excel and UDT macro add-in.

### Transplantation of HSPCs into NCG mice

The transplantation of edited and non-edited HSPCs into mice was adapted from the procedure documented in ^11^. Briefly, frozen aliquots of mobilized peripheral blood CD34+ cells from healthy donors obtained 2 days after editing (D4) were sent to TRANSCURE bioServices (Archamps, France) for xenotransplantation into 4-weeks-old female NODPrkdcem26Cd52 Il2rgem26Cd22/NjuCrl (NCG) mice. Pre-transplant conditioning was based on Busulfan (Sigma, #B1170000) according to the TRANSCURE bioServices protocol. NCG mice followed an acclimatation period of 7 days prior being used in experiments and were housed at a constant temperature T=22 ± 2°C, at a constant relative humidity RH=55 ± 10% and a photoperiod of 12:12-hour light-dark cycle 7am:7pm. A total of 0.75 × 10^6^ edited or non-edited control HSPCs were transplanted via tail vein injection. Sixteen weeks after transplantation, mice were sacrificed, and peripheral blood and bone marrow were recovered. One hundred thousand cells from each organ were harvested, and chimerism was assessed by flow cytometry using the following antibodies: mouse CD45 perCPvio700, Clone# REA747(Miltenyi, #130-110-636), human CD45 BV650, Clone# HI30 (BD, #563717) and viability dye FVS780, (BD, #565388). Editing was also assessed by cytometry using the following antibodies : HLA-E-ABC Vioblue, Clone#REA230 (Miltenyi, 130-120-435), HLA-E APC, Clone1031 (Miltenyi, 130-117-402. The gating strategy used to analyze cellular suspension is documented in supplementary figure 7. All procedures and animal housing for NCG xenotransplantations were performed at TransCure bioServices (Archamps, France) and were reviewed and approved by the local ethics committee (CELEAG).

### Statistical Analysis

Comparisons of numerical variables between two groups were carried out using Mann–Whitney U tests as specified in the figure legends. Two-way ANOVA with Bonferroni post-tests were used to analyze experiments comparing two variables. P-values are indicated in the plots. All statistical analyses were performed using GraphPad Prism v.9.4 (GraphPad).

### Instrument and software used for data acquisition and analysis

The instruments used for this study were similar to the ones used in^11^. Cell viability was acquired using NucleoCounter® NC-250 (ChemoMetec) and analyzed using NucleoView™ software v4.3. CFU colonies were detected using a STEMvision apparatus (STEMCELL Technologies) and analyzed using STEMvision Colony marker (STEMCELL Technologies) v2.0.3.0. DNA quantification was performed using a NanodropOne device (ThermoScientific) and analyzed using NanoDrop QC software v1.6.198. Flow cytometry was performed using Novocyte Quanteon 4025 (Agilent) or Attune NxT (Life Technologies) flow cytometers. BD FACS DIVA software v9.0 (BD), FlowJo v10.8.1 (Treestar), were used to analyze flow cytometry dataset. Single cell formulations were generated using Chromium single cell system (10X Genomics), high throughput DNA sequencing was performed using NovaSeq systems (Illumina). Sequence read were demultiplexed and aligned to the human reference genome (GRCh38), using the CellRanger pipeline v6.1.2 (10X Genomics). GraphPad Prism software v.9.4. was used to plot most of the dataset illustrated in this study.

## V Resources availability

The authors declare that the data supporting the findings of this study are available in the article and in the supplementary information files.

## VI Acknowledgements

Figures include some illustrations created using Biorender.com. Gil Letort, Aymeric Duclert, Diane Le Clerre, Isabelle Chion-Sotinel, Emilie Dessez, Margaux Sevin, Marco Rotondi, Philippe Duchateau & Julien Valton perceived salary from Cellectis S.A. Cellectis S.A funded this research project. Cosimo Ducani & Roger Salvatori perceived Salary from Moligo Technologies.

This paper is dedicated to the memory of our friend and colleague Arianna Moiani, a talented scientist who was part of the early design of this study and unfortunately deceased in 2024 after her courageous battle against cancer.

## VII Author information

### Authors and Affiliations

#### Cellectis S.A., 8 Rue de la Croix Jarry, Paris, France

Gil Letort, Aymeric Duclert, Diane Le Clerre, Isabelle Chion-Sotinel, Emilie Dessez, Margaux Sevin, Marco Rotondi, Philippe Duchateau & Julien Valton

#### Moligo Technologies, Banvaktsvägen 20, 171 48, Solna, Sweden

Cosimo Ducani & Roger Salvatori

## Contributions

P.D., and J.V. conceived the study. G.L. designed, performed and analyzed most of the experiments using HSPCs as starting biological material including the preparation of cellular sample for CITE-Seq analysis. A.D. analyzed all CITE-Seq datasets with the help of G.L. and J.V., who participated in curating and plotting datasets. DL.C., I.C-S. and M.S. participated to the establishment of the optimal experimental conditions of HSPCs engineering. E.D. performed and analyzed most of the experiments using T-cells as starting biological material and M.R., supervised E.D., experimental work. R.S., produced the LssDNA and CssDNA used in this study and C.D., supervised R.S., experimental work and the collaboration between Moligo Technologies and Cellectis. SA. G.L., DL.C., I.C-S., M.S. and J.V., designed the in vivo study performed with NCG mice and G.L., I.C-S. analyzed the dataset. J.V. supervised the study and wrote the manuscript with the help from all authors. J.V. and C.D., established and supervised the collaboration between Cellectis and Moligo Technologies that was coordinated with the help of M.S.

