## Supplementary materials for "Circularization of Single-Stranded DNA Donor Template Unleashes the Power of Non-Viral Gene Delivery for Long-Term HSCs editing"

1. Cellectis S.A., 8 Rue de la Croix Jarry, 75013 Paris, France.

2. Moligo Technologies, Anderstorpsvägen 16, 171 54, Solna, Sweden

3. These authors contributed equally

4. Lead contact

**Supplementary information**

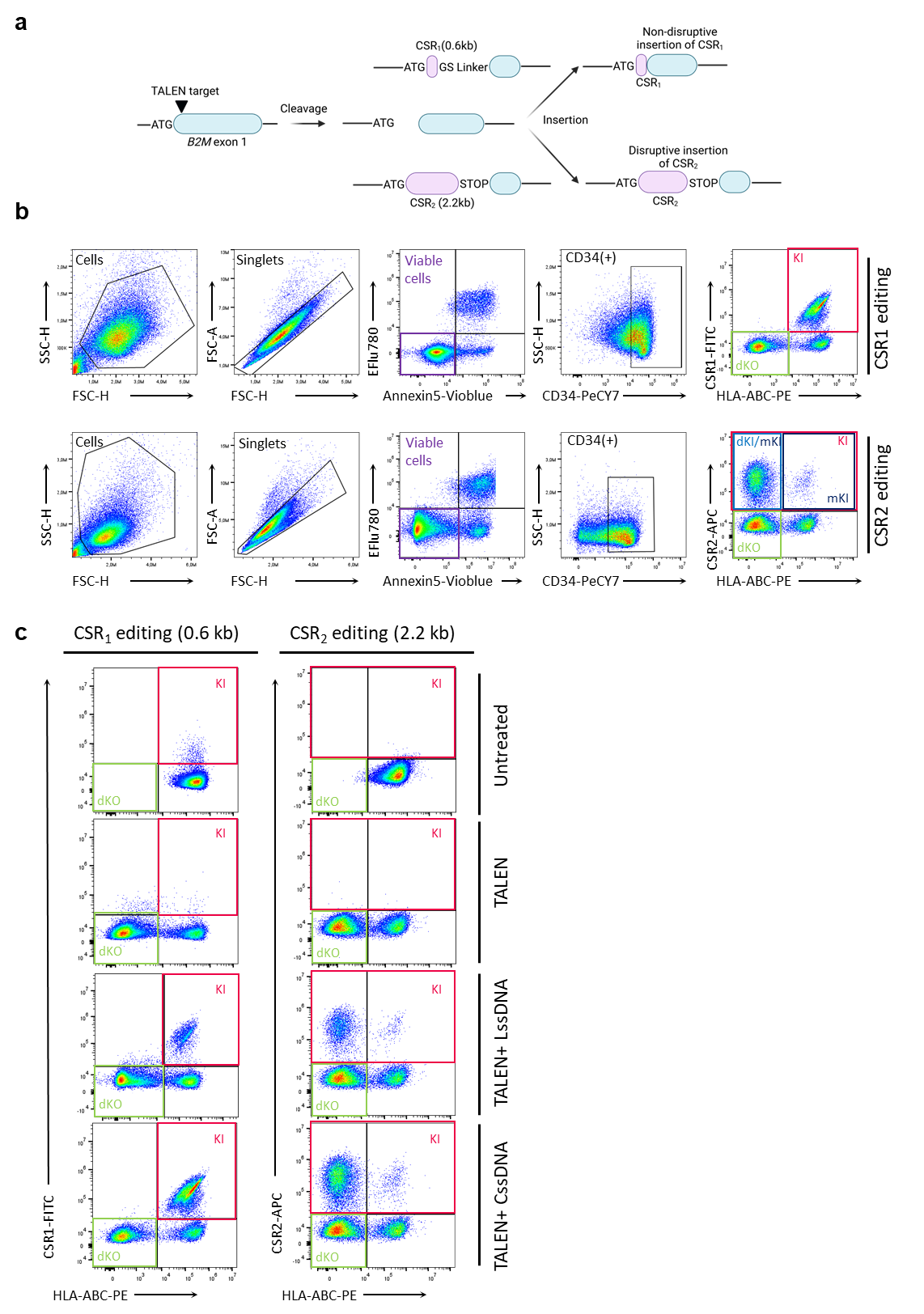

**Supplementary figure 1. Flow cytometry analysis of HSPCs edited with TALEN and CssDNA or LssDNA. a,** Scheme illustrating the TALEN gene editing strategy used to insert CSR_1_ (0.6kb) and CSR_2_ (2.2kb) DNA donor templates at the B2M exon 1 locus in a non-disruptive and disruptive manner, respectively. **b,** FCM gating strategy to assess the frequency of viable CD34 (+) cells harboring mono and double allelic KI (mKI, dKI and KI, respectively) and double KO (dKO) events in HSPCs edited with TALEN-B2M and ssDNA encoding CSR_1_ and CSR_2_. The Ratio KI/KO documented in main figures 1, 2 and supplementary figure 4 are obtained by computing KI/dKO for CSR1 construct and (mKI+dKI)/dKO for CSR2 construct. **c,** Representative FCM results obtained with HSPCs either unedited or edited in presence of TALEN or TALEN and CssDNA encoding CSR_1_ and CSR_2_.

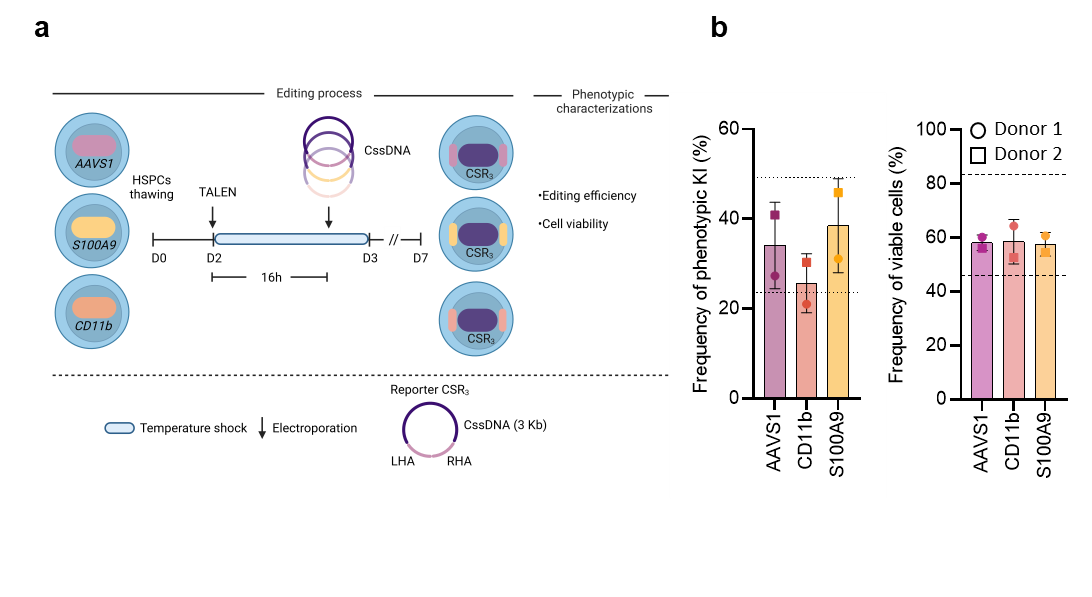

**Supplementary figure 2. CssDNA and TALEN enable efficient Knock-in at multiple genes in HSPCs. a.** Representative schema of HSPCs editing protocol using an mRNA encoding TALEN targeting the *AAVS1, CD11B or S100A9* loci and cssDNA as DNA donor templates to insert a reported gene (CSR_3_) via non-disruptive. **b.** Experimental results obtained on 2 independent biological donors, illustrating the frequency of cells harboring phenotypic KI events and their viability measure at D7 (n=2 independent HSPC donors). The dashed lines illustrate the range of KI frequency and cell viability obtained with HSPCs edited by the TALEN targeting B2M and the CssDNA encoding the CSR_2_ (2.2kb).

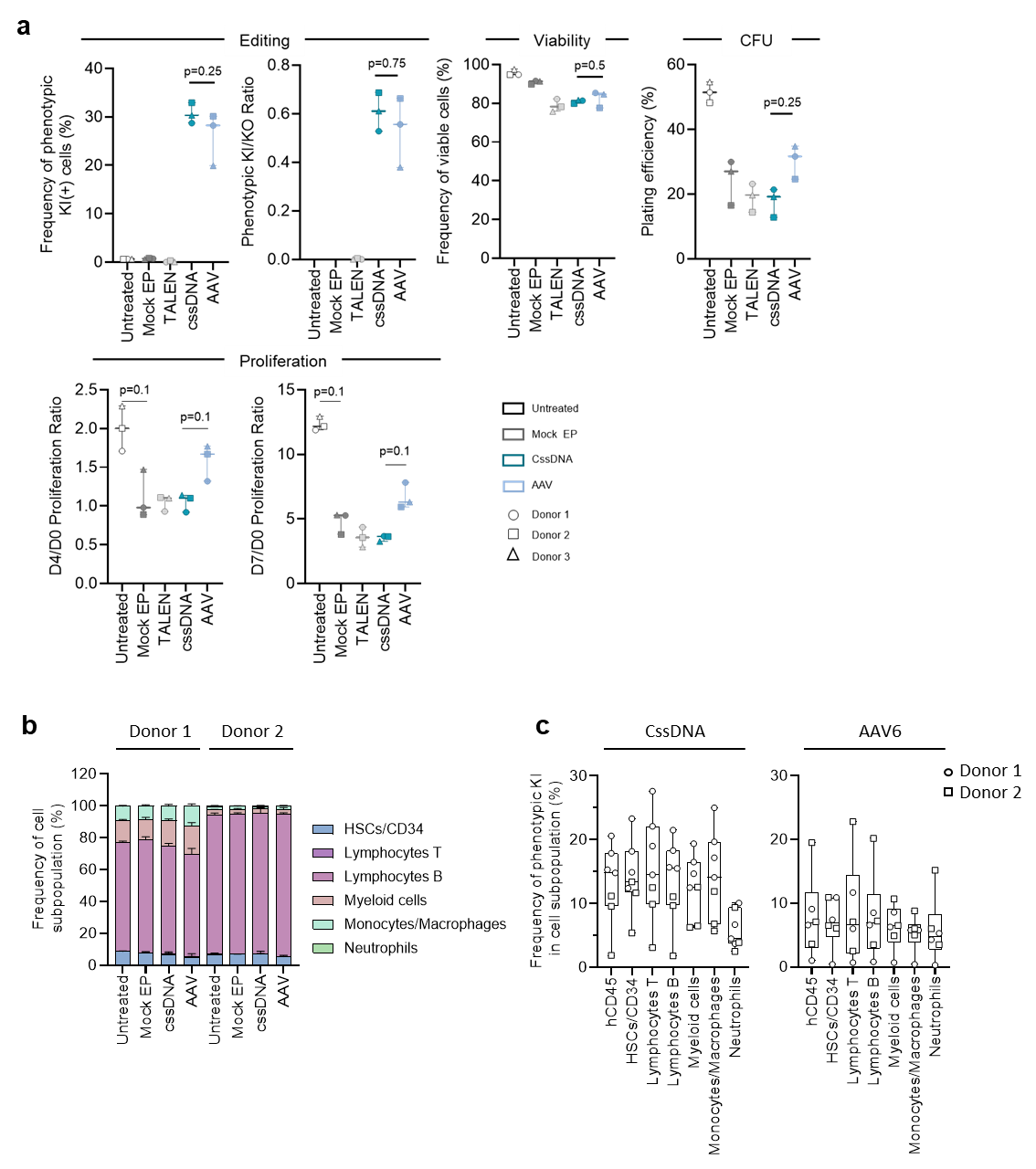

**Supplementary figure 3. CssDNA and TALEN-mediated knock-in process leads to a higher engraftment of KI(+) HSPCs compared to the AAV and TALEN reference process**. **a**, in vitro results illustrating the frequency of KI, the ratio of KI/KO, the viability, differentiation and proliferation capacities of HSPC edited by TALEN in the presence of CssDNA or AAV6 (n=3 independent HSPC donors). **b**, *in vivo* differentiation capacity of edited HSPC characterized in the bone marrow of NCG mice, 16 weeks post HSPC injection onset. **c**, Frequency of KI obtained in different cell subpopulations detected in NCG mice, 16 weeks post HSPC injection onset (n=2 independent HSPC donors).

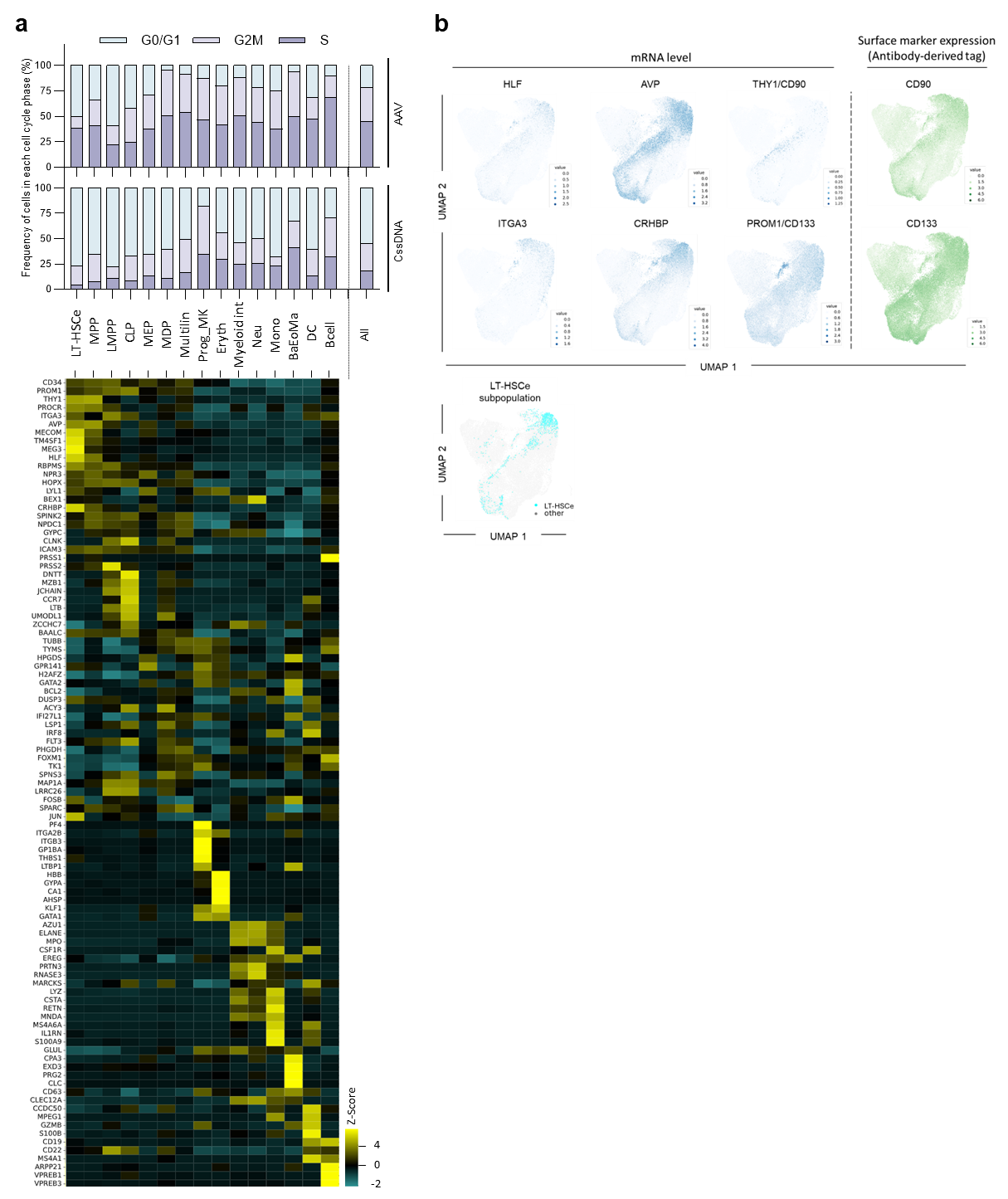

**Supplementary figure 4. Identification of Long-Term HSC enriched cell (LT-HSCe) subpopulation by CITE-Seq analysis of non-edited and edited HSPCs at D4. a top panel**, Frequency of cells identified in G0/G1, G2M and S cell cycle phase among each cellular subpupolation in UMAP (n= 3 donors). **a bottom panel**, Hitmap showing Z-score of gene expression in each subpopulation identified in UMAP (n= 3 donor)**. b, Top panel,** UMAP illustrating the level of HLF, AVP, ITGA3 and CRHBP mRNA transcripts and CD90 (THY1) and CD133 (PROM1) surface marker expression determined by CITE-Seq. **b, bottom panel**, UMAP illustrating the position of LT-HSCe subpopulation identified by cell typing using^22^. **a bottom panel and b** were obtained out of 3 independent HSPC donors, by aggregating untreated, CssDNA and AAV experimental conditions, representing 22359 encapsulated cells.

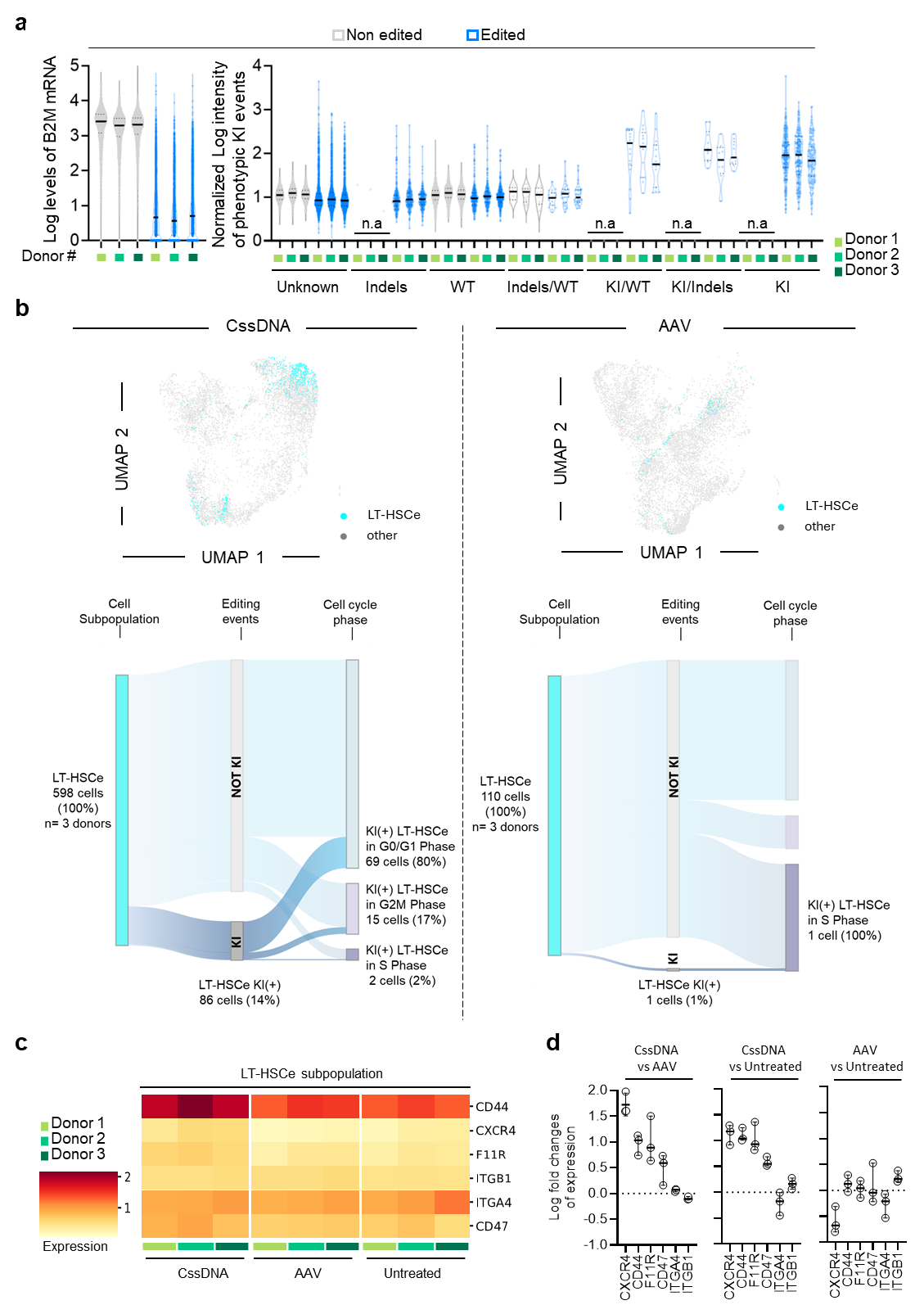

**Supplementary figure 5. CITE-Seq analysis of non-edited and edited HSPCs at D4. a. left panel,** levels of B2M transcripts determined by CITE-Seq in non-edited HPSCs and HSPCs edited with TALEN and DNA donor template. **a. right panel,** Normalized intensity of the surface expression of CSR_2_ (phenotypic KI events) as a function of different genotypes (WT, wild type; Indels, NHEJ-mediated insertion and deletion and KI, CSR_2_ insertion) identified at the mRNA level (n.a. indicates dataset with less than 2 cells detected). **b, upper panel,** UMAP showing the position of LT-HSCe subpopulation identified in CssDNA- and AAV-edited HSPCs. **b. lower panel,** Sankey diagram illustrating the frequency of phenotypic KI(+) and KI(-) LT-HSCe in G0/G1, G2M and S phases identified in *in vitro* CssDNA- and AAV-edited HSPCs experimental group at D4, right before injection in NCG murine model. **c.** heatmap illustrating the relative expression of cell surface markers known to be involved in bone marrow niche adhesion, in the CssDNA- and AAV-edited and untreated HSPCs. **d.** Log fold change of expression of cell surface markers in the CssDNA-edited group versus the AAV-edited or untreated groups, and in the AAV versus untreated groups, obtained at D4, right before injection in NCG murine model (n=3 independent HSPC donors).

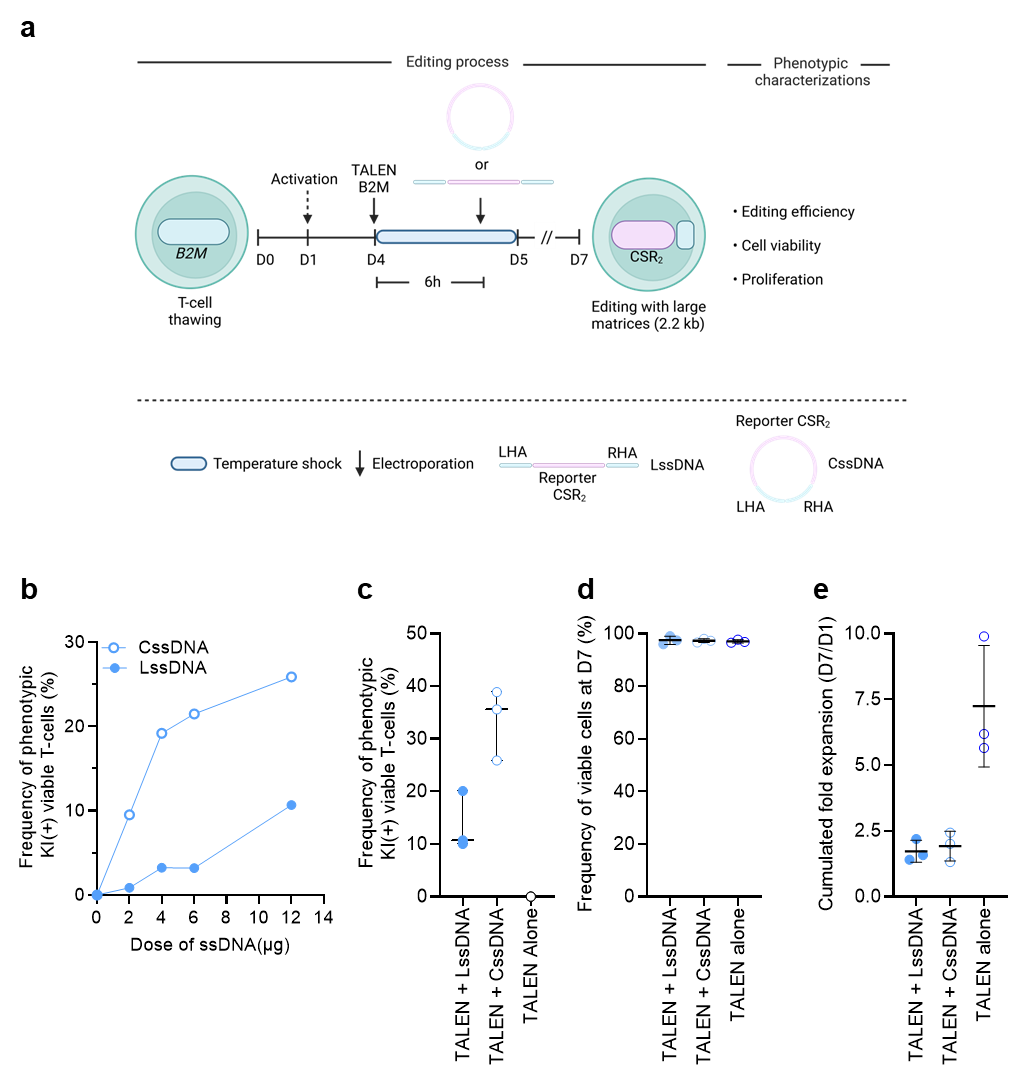

**Supplementary figure 6. Circularization of ssDNA increases the overall efficiency of TALEN-mediated Knock-in in primary T-cells. a.** Representative schema of T-cell editing protocol using an mRNA encoding TALEN targeting the *B2M* locus and LssDNA or CssDNA as DNA donor templates to insert a CSR_2_ (2.2 kb) reported gene via disruptive insertions. **b.** Frequency of KI obtained by FCM as a function of ssDNA dose**. c.** Frequency of phenotypic KI obtained by FCM using TALEN_B2M_ and 12 µg of LssDNA or CssDNA at D7. **d.** Viability of T-cells edited by TALEN_B2M_ and LssDNA or CssDNA and by TALEN_B2M_ alone at D7. **e.** Proliferation capacity of edited cells illustrated by their cumulated fold expansion between D1 and D7 (n=3 independent T-cell donors).

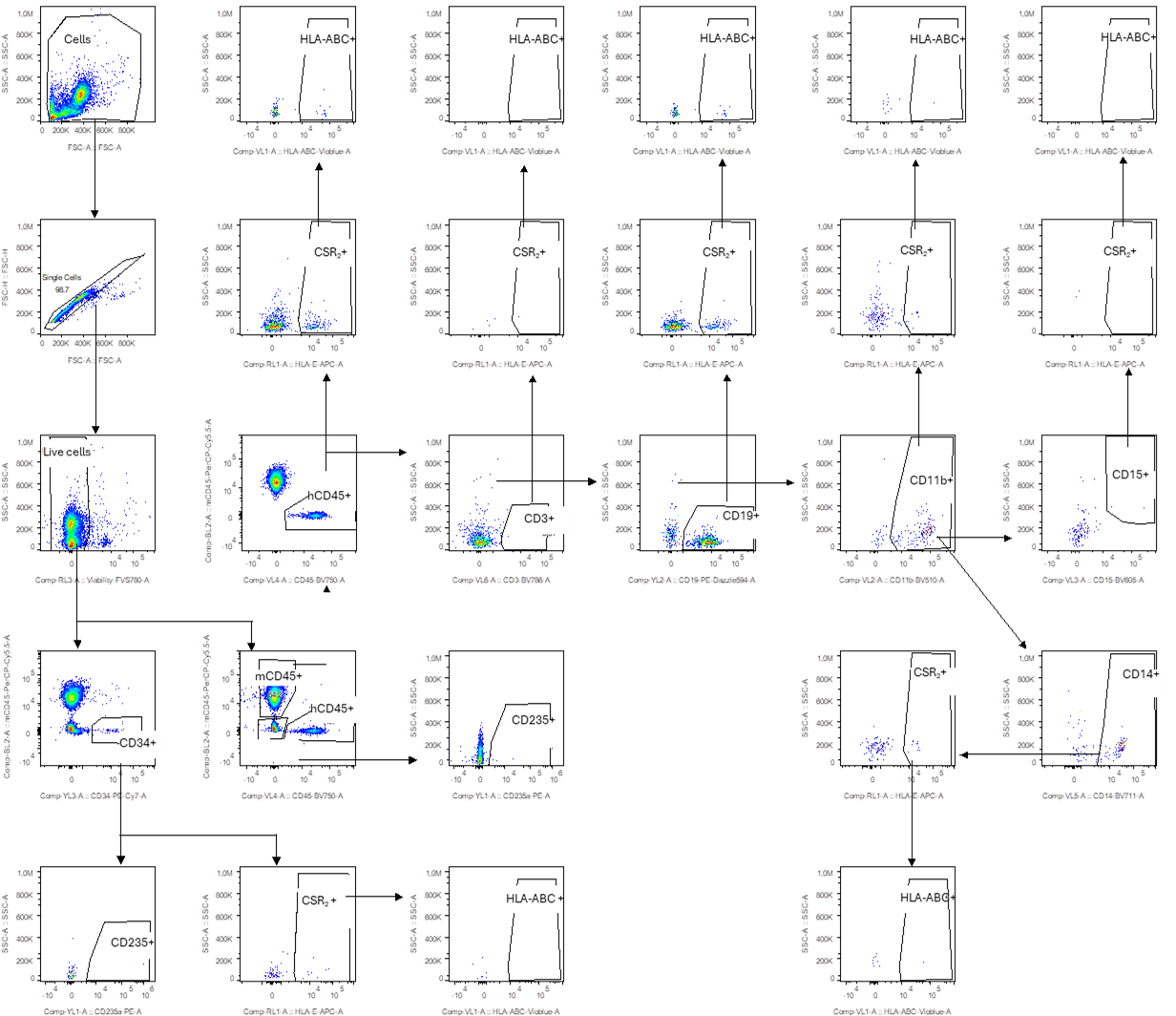

**Supplementary figure 7. Gating strategy to assess the efficiency of edited HSPCs engraftment in NCG mice.** Representative gating strategy used to determine the level of hCD45 engraftment and the level of phenotypic KI and KO event in different tissues of NCG mice, 16 weeks post edited HSPCs injection.

**Table**

**Supplementary Table 1.** CITE-Seq dataset specifications

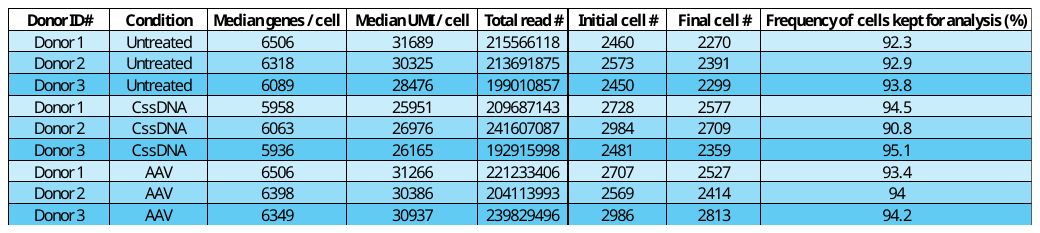

**Supplementary Table 2.** Antibody-related tags used for CITE-Seq procedure.

|  | **Catalog** | **Description** | **Clone** | **Barcode** |
| --- | --- | --- | --- | --- |
| **CITE-Seq Antibodies** | 399905 | TotalSeq™-C Human Universal Cocktail, V1.0 | - | Multiple |
|  | 341507 | CCR10 | - | ATCTGTATGTCACAG |
|  | 323309 | CD109 | - | CACTTAACTCTGGGT |
|  | 362011 | CD130 | - | CACGAGAATTTCAGT |
|  | 372817 | CD133 | - | TGGTAACGACAGTCC |
|  | 324813 | CD164 | - | GAGGCACTTAACATA |
|  | 329229 | CD200 | - | CACGTAGACCTTTGC |
|  | 351909 | CD201 | - | GTTTCCTTGACCAAG |
|  | 308217 | CD253 | - | GCCATTCCTGCCTAA |
|  | 307415 | CD262 | - | TTGGCCTGTAGAAAT |
|  | 343541 | CD34 | - | GCAGAAATCTCCCTT |
|  | 345049 | CD366 | - | TGTCCTACCCAACTT |
|  | 342325 | CD66a/c/e | - | GGGACAGTTCGTTTC |
|  | 326813 | CD74 | - | CTGTAGCATTTCCCT |
|  | 328145 | CD90 | - | GCATTGTACGATTCA |
|  | 315607 | CD98 | - | GCACCAACAGCCATT |
|  | 373209 | HLA-F | - | GCAACTCTCCTACCT |
|  | 361113 | MR1 | - | GTTGGATGGTAGACT |
|  | 901535 | HA-tag | - | TTGTGAGGCTTCTCT |
| **Hashtag** | 394661 | CD298/β2-microglobulin | HT-1 | GTCAACTCTTTAGCG |
|  | 394663 | CD298/β2-microglobulin | HT-2 | TGATGGCCTATTGGG |
|  | 394665 | CD298/β2-microglobulin | HT-3 | TTCCGCCTCTCTTTG |
|  | 394667 | CD298/β2-microglobulin | HT-4 | AGTAAGTTCAGCGTA |
